# Modulation of functional phosphorylation sites by basic residues in the Unique domain of c-Src

**DOI:** 10.1101/2023.05.23.541872

**Authors:** Andras Lang, Alejandro Fernández, Mireia Diaz-Lobo, Mar Vilanova, Francisco Cárdenas, Margarida Gairí, Miquel Pons

## Abstract

In contrast to the well-studied canonical regulatory mechanisms, the way by which the recently discovered Src N-terminal regulatory element (SNRE) modulates Src activity is not yet well understood. Phosphorylation of serine and threonine residues modulate the charge distribution along the disordered region of the SNRE and may affect a fuzzy complex with the SH3 domain that is believed to act as an information transduction element. The preexisting positively charged sites can interact with the newly introduced phosphate groups by modulating their acidity, introducing local conformational restrictions, or coupling various phosphosites into a functional unit. In this paper we use pH dependent NMR measurements combined with single point mutations to identify the interactions of basic residues with physiologically important phosphorylated residues and to characterize the effect of these interactions in neighbor residues, thus providing insight on the electrostatic network in the isolated disordered regions and in the entire SNRE. From a methodological point of view, the linear relationship observed between the mutation induced pKa changes of the phosphate groups of phosphoserine and phosphothreonine and the pH induced chemical shifts of the NH groups of these residues provides a very convenient alternative to identify interacting phosphate groups without the need to introduce point mutations on specific basic residues.

## 1. Introduction

c-Src is the leading member of the Src family of kinases (SFK) and the first discovered proto-oncogene. It mediates signal transduction processes related to cell migration, invasion, and survival [1]. High levels of c-Src activity are associated to poor prognosis in different cancers [2,3]. All SFK share the same domain architecture formed by highly conserved globular Src homology domains (SH1, SH2, and SH3) and an N-terminal disordered region formed by the SH4 and Unique domains. The canonical regulatory mechanisms involving the globular domains have been extensively studied [4,5]. However, the regulation layer involving the intrinsically disordered region of c_Src [6, 7], remains poorly understood. Phosphorylation of serine and threonine residues are important players in the regulation by disordered domains [8]. Dephosphorylated serine 17 (S17), favors Src dimerization and is found preferentially in cancer cells [9, 10]. Phosphorylation of serine 75 (S75) causes changes in cell growth, cytoskeletal reorganization and mediates ubiquitination and degradation [11,12]. Phosphorylation of threonine 37 (T37) activates Src by disrupting the interaction between the SH2 domain and regulatory phosphotyrosine 530 [13]. Phosphorylation of serine 43 (S43) and serine 51 (S51) by Wnt3A have opposite effects on Src activation [14].

Understanding the distinct and specific effects of phosphorylation at each of the sites starts by determining the functional unit that is affected by the introduction of the phosphate group. Is it strictly the modified residue or there are neighbor residues specifically affected, thus defining an extended functional site? Basic residues are obvious candidates to extend the functional effects of phosphorylation through interaction of the positively charged side chains with the negatively charged phosphate. Interestingly, in the Unique domain of c-Src, five of the functionally important serine and threonine phosphorylation sites have basic residues located three or less residues away along the sequence: S17 follows a stretch of three arginine residues, S75 is three residues away from arginine 78 (R78), lysine 40 (K40) is located between threonine 37 (T37) and serine 43 (S43) and S51 is three residues away from an arginine residue.

To test the formation of specific contacts between phosphorylated serine or threonine residues and proximal basic groups we have measured the changes in phosphate pKa caused the mutation of individual basic groups. We show that the mere presence of a basic group in the proximity of the phosphate group does not imply the two groups are interacting. Additionally, by observing the effect of pH on the NH residues of wild-type phosphorylated c-Src we can identify the ensemble of residues that are directly or indirectly affected by the titration of the phosphorylated residues. By comparing the pH dependency of NH chemical shifts in wild type c-Src with the results obtained with mutated forms we demonstrate a very simple NMR method to identify the phosphorylated residues that are interacting with basic residues, that does not require mutagenesis or complete pKa determination.

The Unique, SH4 and SH3 domains form the Src N-terminal regulatory element (SNRE) characterized by a fuzzy complex in which the Unique and SH4 domain form multiple, weak, dynamic contacts with the SH3 domain, while remaining disordered. The SH3 domain couples with the other globular domains presumably transmitting signaling events, such as phosphorylation of the disordered region to the kinase domain [15,16]. Electrostatic interactions are an important component of the fuzzy complex that can be modulated by phosphorylation [17]. The number and distribution of charged residues has been recognized as an important parameter affecting the conformational ensemble sampled by disordered protein regions [18]. Not surprisingly, phosphorylation of the Unique domain affects the fuzzy complex and the pKa of the phosphorylated residues are sensitive probes of this interaction.

We have centered the study on the sites that are phosphorylated by the mitogen activated kinase ERK2, a promiscuous kinase that preferentially phosphorylates serine or threonine residues followed by proline [19]. Unexpectedly, we found that ERK2 phosphorylates not only T37 and S75, but also S43, and to a minor extent, S51 although S43 and S51 are not followed by proline.

## 2. Results

### 2.1. Phosphorylation of the Unique domain of c-Src by ERK2

The main phosphorylation sites in the Unique domain of human c-Src are T37 and S75, both of which are followed by proline residues. The S/TP motif is the consensus sequence for mitogen activated kinases. We chose ERK2 for the in vitro phosphorylation of these two residues because of the availability of a constitutively activated ERK2 form [20]. Surprisingly, native gel electrophoresis of USH3 treated with ATP and ERK2 gave three major bands corresponding to the incorporation of up to three phosphate groups (Figure 1), suggesting the presence of at least an additional ERK2 target site, besides T37 and S75.

**Figure 1.**
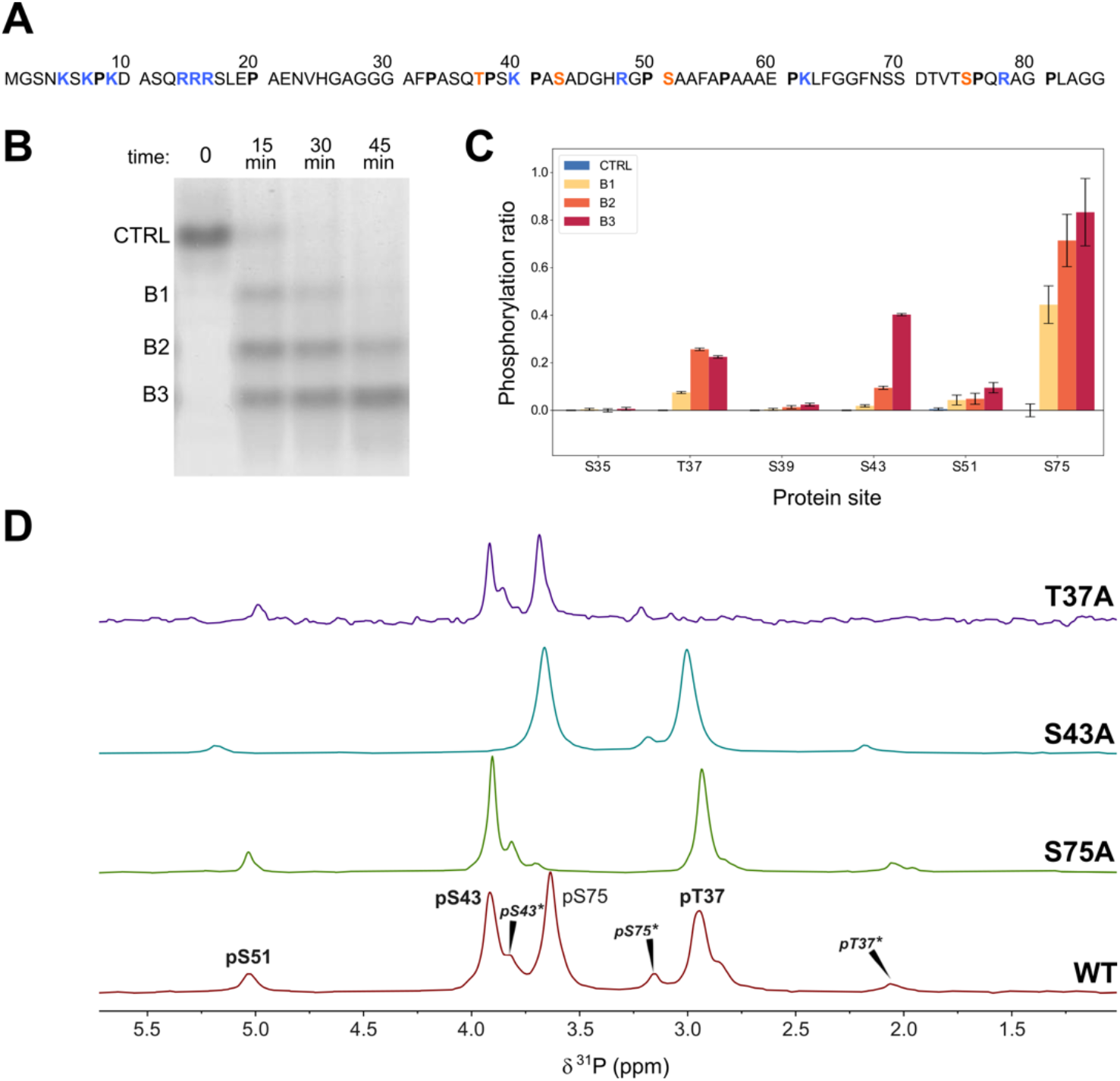
A. Sequence of the disordered region of human c-Src with the phosphorylated residues studied in this work highlighted in red, basic residues marked in blue, and proline residues in bold face. B. Native polyacrylamide electrophoresis during the phosphorylation of USH3 with ERK2. Bands are separated by the number of phosphate groups incorporated. C. Phosphorylated sites detected by MS/MS in the three electrophoretic bands corresponding to the global incorporation of one, two and three phosphates in USH3. The pS75 site is located in a different peptide than the other sites. D. ^31^P-NMR (164 MHz) of wild-type USH3 and mutants used for NMR assignment. Asterisks denote peaks from forms containing cis proline bonds. The spectra were recorded in 50 mM 2-(N-morpholino)ethanesulfonic acid (MES) pH 6.3 and referenced to triethylphosphate at 0,44 ppm.

Tandem mass spectrometry analysis of the tryptic peptide that contains the T37 site confirmed that S43 was extensively phosphorylated by ERK2. An additional minor phosphorylation site, S51 was detected by mass-spectrometry (Figure 1). These sites do not contain a canonical ERK2 substrate motif. Phosphorylation of S43 was confirmed by ^31^P-NMR (*vide infra*). Minor signals assigned to phosphorylated S51 were also detected by ^31^P-NMR and in some ^1^H-^15^N correlation spectra, and confirmed by sequential NMR assignment in ^15^N,^13^C labeled samples.

### 2.2. ^31^P-NMR

Figure 1 shows ^31^P-NMR spectra of triple phosphorylated USH3 at pH 6.3 and 298K. The peaks of pS75 and pS43 overlap at pH 7. The three main peaks were assigned by comparison with mutated constructs in which the individual serine and threonine phosphosites were replaced by alanine. The disappearance of the signal at 3.0 ppm in the S43A mutant confirmed the phosphorylation of S43 by ERK2. Additional low intensity peaks disappeared in each of the mutants, suggesting that they arise from the coexistence of phosphorylated peptides with a cis proline bond. The presence of cis proline forms was confirmed in ^13^C and ^15^N enriched samples (*vide infra*).

pT37 and pS75 that are located next to proline residues showed the largest upfield shifts when the proline adopts a cis conformation (ca. 0.9 ppm for T37 and 0.5 ppm for S75). pS43, which does have a direct proline neighbor show two minor forms at 0.1 and 0.2 ppm upfield, with relative integrals 0.1 and 0.01 that suggests that pS43 is experiencing the effect of two different cis proline bonds.

The minor signal at around 4 ppm that is present in all the mutants, was tentatively assigned to the small amount of phosphorylated serine 51 detected by mass spectrometry. Consistent with this assignment, the chemical shift of this minor signal is most affected by the S43A mutation, which is the one closest to S51.

The pH dependence of the ^31^P-NMR signals was used to determine the apparent pKa values of the individual phosphate groups in the triple phosphorylated form of USH3 at 298K. The relevant pKa values were extracted by non-linear fitting of the Henderson-Hasselbalch equation. The pKa values of pT37, pS43 and pS75 are 6.10 ± 0.01, 5.80 ± 0.01, and 6.03 ± 0.01 respectively (Table1).

**Table 1.** pKa values in wild type USH3, USrc and basic residue mutants^1^.

| Phosphosite<sup>1</sup> | USrc<br>wild type | USrc<br>K40A | USrc<br>R48A<sup>3</sup> | USrc<br>R78A | USH3<br>wild type |
| --- | --- | --- | --- | --- | --- |
| pT37 | 6.19 | 6.18 (-0.01)<sup>2</sup> | 6.09 (-0.10) | 6.16 (-0.03) | 6.10 (-0.09) |
| pS43 | 5.88 | 5.98 **(+0.10)** | - | 5.85 (-0.03) | 5.80 (-0.08) |
| pS75 | 6.06 | 6.01 (-0.05) | 5.96 (-0.10) | 6.26 **(+0.20)** | 6.03 (-0.03) |

<sup>1</sup> <sup>31</sup>P-NMR; T=298 K.<sup>2</sup> Values in parentheses are the pKa difference with respect to wild type phosphorylated USrc. Error limits estimated from Monte-Carlo analysis are ± 0.01 (see experimental part)<sup>3</sup> Phosphorylated only at T37 and S75

The pKa of the triple phosphorylated protein was also measured in USrc (Figure 2). The pKa values of pT37 and pS43 in the absence of the SH3 domain are 6.19 ± 0.01 and 5.88 ± 0.01, respectively, an increase of around 0.08 units. The pKa of pS75 shows a smaller increase of only 0.03 units, although S75 is closer in the sequence to the SH3 domain. This observation is compatible with the hypothesis that the pKa differences in the presence and in the absence of the SH3 domain are caused by changes in the fuzzy complex.

**Figure 2.**
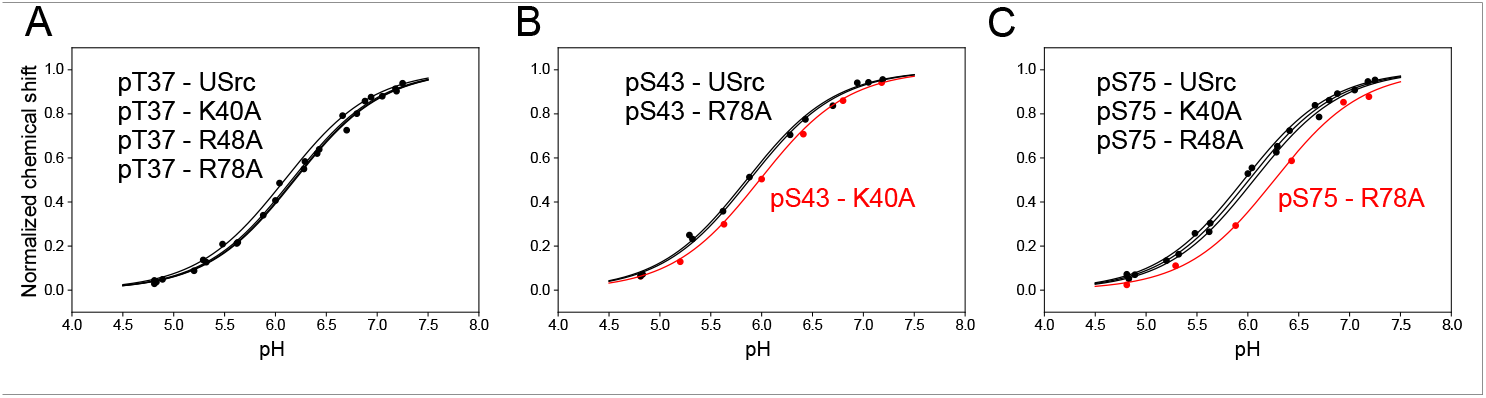
Normalized ^31^P-NMR chemical shift changes of (A) pT37, (B) pS43 and (C) pS75 phosphoresidues in USrc and single basic residue mutants. pKa shifts indicating the release of the interaction between pS43 and K40, and of pS75 with R78 are highlighted in red.

To study the effect of the basic residues, we measured the pKa values of phosphate groups when residues K40, R48 and R78 were individually mutated to alanine in USrc (Figure 2). Phosphorylation of the R48A mutant produced only a double phosphorylated species containing pT37 and pS75, with S43 not phosphorylated.

Replacing the positively charged K40 by alanine causes a shift of +0.10 units in the pKa of pS43 (Table1) strongly suggesting the formation of a salt-bridge between the side chain of K40 and the phosphate group of pS43 that increases its acidity. Interestingly, the K40A mutation has no effect on the pKa of pT37, although the distances of K40 to T37 and S43 along the sequence are the same. The very different effect of the K40A mutation on these phosphate groups strongly suggests a direct interaction between the side chains of K40 and the phosphate group of pS43, as the electrostatic field generated by the positive charge in the lysine side chain is expected to be sensed similarly by the phosphate groups of pT37 and pS43.

The pKa of pS75 experiences a large shift (+0.20 units) when R78 is mutated to alanine, also suggesting the formation of a salt-bridge between the arginine side chain and the phosphate group.

The K40A mutation has a significant (−0.05 units) long-range effect on pS75 although in the opposite direction to that caused in pS43. This is an indirect effect probably caused by changes in the average distance of pS75 to the region including T37, K40, and S43. Long distance effects were observed in unphosphorylated USrc by paramagnetic relaxation experiments [21]. These long-range interactions also exist in the fuzzy complex with the SH3 domain.

### 2.3. ^1^H-^15^N-HSQC

Phosphorylation of serine and threonine residues cause a very large shift in the NH peak of the phosphorylated residue. ^1^H-^15^N-HSQC experiments of USH3 and USrc phosphorylated with ERK2 show three main peaks in the region corresponding to phosphorylated serine and threonine, all of which showing a strong pH dependency Figure 3). This contrasts with previous reports showing that only T37 and S75 were phosphorylated by CDK5/p25 [21]. The new peak was assigned to pS43, confirming the mass spectrometry results.

**Figure 3.**
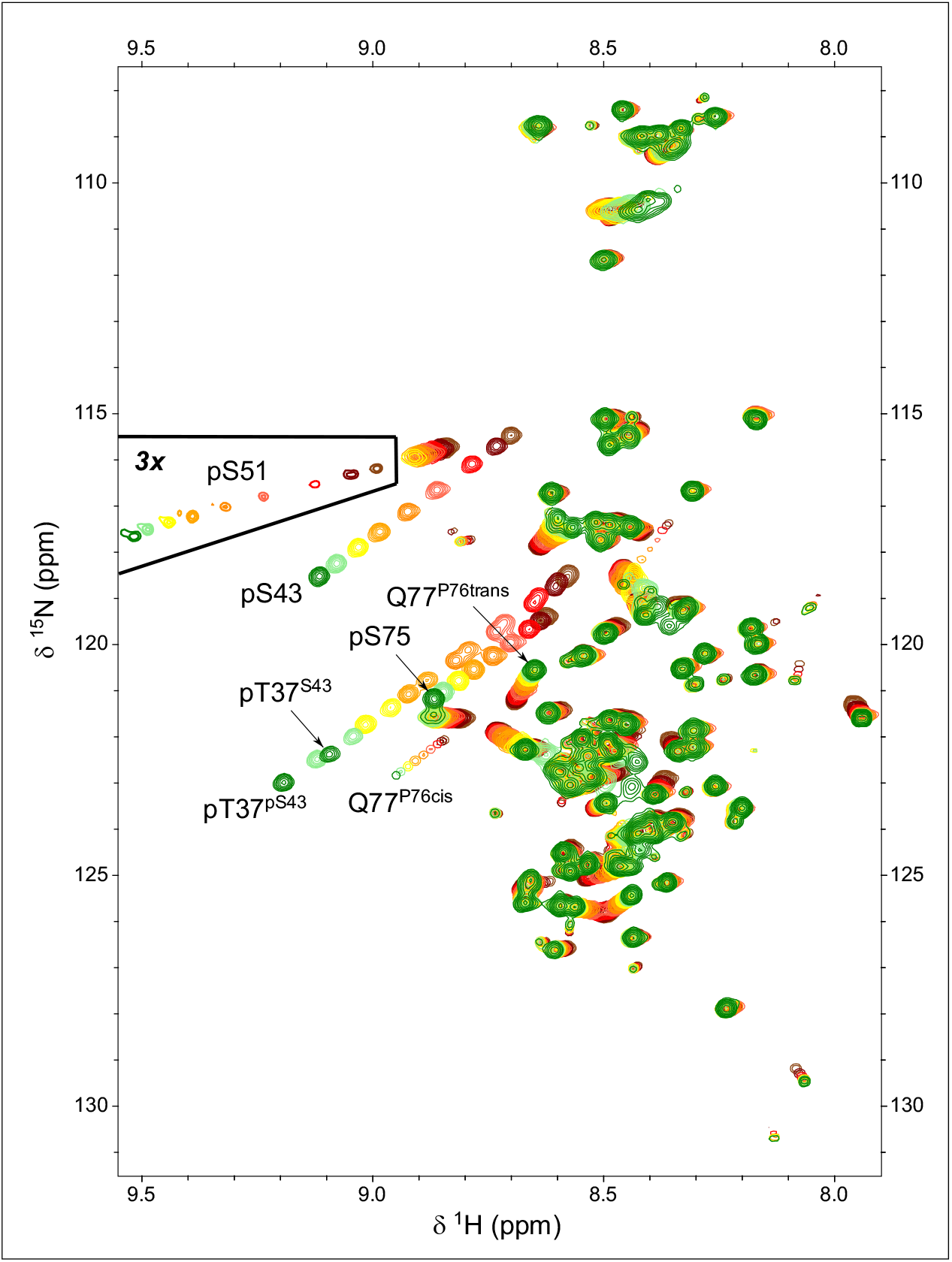
pH dependency of ^1^H-^15^N-HSQC spectra of ERK2-phosphorylated USrc between pH 7.2 (green) and pH 4.7 (brown). The peaks showing the largest pH dependency are those of the backbone NH of the phosphorylated residues. The marked region was plotted at a lower contour level to show the small quantity of phosphorylated S51 present in this sample.

The NH signals from the three phosphorylated residues, pT37, pS43, and pS75 shift downfield significantly in the proton and nitrogen dimensions with a very similar slope on going from the predominantly single protonated (i.*e*. monoanionic phosphate) form at pH 4.7 to the unprotonated (*i*.*e*. dianionic phosphate) form at pH 7.2. Thus, the corresponding peaks move following parallel lines (Figure 3).

A weak signal in the HSQC spectrum of USrc could be unequivocally assigned to the small amount of phosphorylated S51 and the pKa of this minor form could be determined to be 5.61±0.01. The phosphate group in pS51 is more acidic than those of the other serine residues in USrc. The pKa of the small signal at low field in the ^31^P-NMR spectra of the K40A mutant USrc, which gave the strongest signal, has an apparent pKa of 5.72, consistent with the tentative assignment of this signal to pS51.

Phosphorylation also affects the NH chemical shifts of residues other than the phosphorylated ones. In those residues, the relative effects of pH on the ^1^H and ^15^N dimensions may be different for those of the phosphorylated residues, thus the peaks follow lines with different slopes. The direction of the peak displacement reflects the combination of various types of interactions such as electrostatic effects, hydrogen bonding, or conformational shifts that may affect differently the two nuclei.

The pKa measured from these signals can be used to identify the titrating group primarily responsible of the observed pH dependency. This is the case of the NHε of the guanidinium side chain of R78 that titrates with a pKa of 5.94 ± 0.01, identical within the experimental error, to that measured in the NH of pS75 (5.93 ± 0.01), confirming the direct interaction between pS75 and R78 side chains. The carboxamide NH_2_ signals and backbone NH of Q77 are also titrated with the same apparent pKa (5.92± 0.02) (Figure 4).

**Figure 4.**
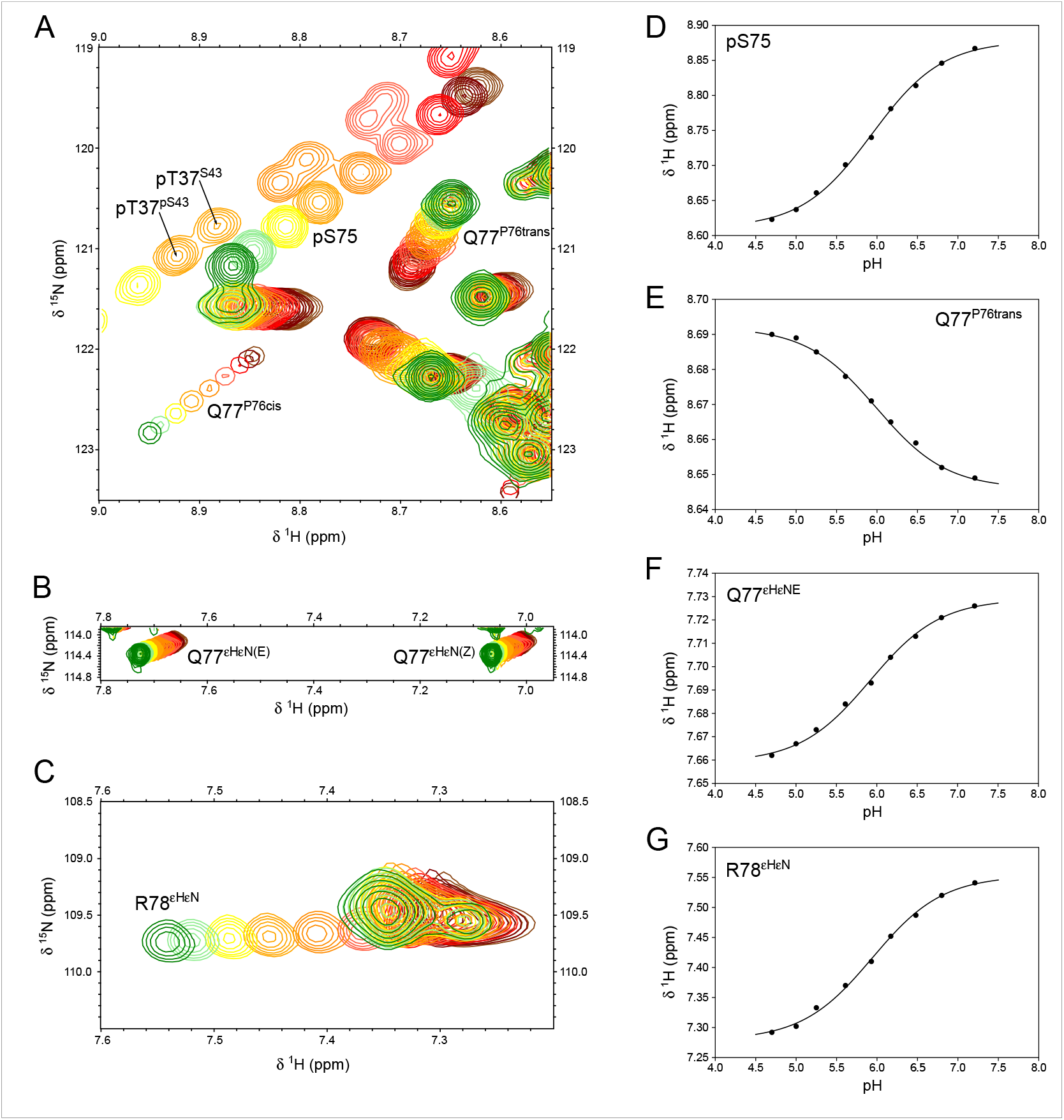
^1^H-^15^N-HSQC-monitored pH titration of ERK2-phosphorylated Usrc. A-C. Expansions of spectra showing the effect of titrating the phosphate groups between pH 7.2 (green) and pH 4.7 (brown). Labeled peaks are those affected by the titration of pS75, except pT37. The R78 side chains shown in C are folded in the ^15^N-dimension. D-G: The various peaks affected by the titration of pS75 show the same pKa value. The identical pKa measured in pS75 backbone NH and the side chains of R78 and Q77 confirm the direct interaction between pS75 and R78. The two sets of peaks for pT37 arise from the coexistence in this sample of species with phosphorylated and unphosphorylated S43 that have distinct pT37 pKa values.

The HSQC experiments reveal additional peaks that could be assigned to residues next to proline in the cis conformation. The configuration of the proline peptide bond could be determined from the difference between the chemical shifts of the β and γ carbons, which is around 10 ppm in the cis form and only 5 ppm in the trans configuration [22]. The pH-induced chemical shifts of Q77 NH in the major trans form go in the opposite direction of those of pS75 (Figure 4D and E) suggesting an indirect effect affecting the Q77 side chain when the phosphate group of pS75 and the guanidinium group of R78 interact.

The NH main chain signal of Q77 in the species with a cis and trans peptide bond at proline 76 are well resolved. The pH-induced chemical shift changes of the NH of Q77 in the cis form follow a similar slope than those of pS75 (Figure 3) suggesting that the interaction between the phosphate and the pS75 and Q77_(P76 cis)_ backbone NH are similar, which is compatible with the proximity of Q77 and S75 when the intervening proline is in the cis form.

Phosphorylation of S43 by ERK2 is slower than that of T37 and S75 that contain the consensus target sequence. Thus, it was possible to prepare samples that contained a mixture of phosphorylated and non-phosphorylated S43 that could be easily distinguished in the HSQC experiments. This allowed to determine the effect of the phosphate group in S43 on the pKa of pS37 (Figure 4A): the presence of the negatively charged pS43 makes the phosphate group of pT37 less acidic by +0.07 units.

### 2.4. Formation of a fuzzy complex affects the pKa of phosphorylated residues

The pKa values of pT37 and pS43 measured at 298K in the presence of the SH3 domain are 0.09 and 0.08 units lower, respectively, than those measured in the isolated disordered region (Table1). The difference is smaller for pS75 (−0.03 units). The differences observed at 278K follow the opposite trend: the pKa values increase when the SH3 domain is present, and the largest changes are observed for pS75 (+0.06) and pS43 (+0.06) and the pKa of pT37 shifts by +0.03 units. (Table2). All the changes are significant and suggest the electrostatic environment of the phosphorylated groups is affected by long-range interactions in the fuzzy complex formed between the disordered and SH3 domains.

**Table 2.** Temperature dependent pKa in the fuzzy complex and isolated disordered domains

| Phosphosite<sup>1</sup> | Temp.<sup>2</sup> | pT37 | pS43 | pS75 |
| --- | --- | --- | --- | --- |
| pUSrc | 278 | 6.13 | 5.80 | 5.93 |
| pUSrc | 298 | 6.19 | 5.88 | 6.06 |
| pUSH3 | 278 | 6.16 | 5.86 | 5.99 |
| pUSH3 | 298 | 6.10 | 5.80 | 6.03 |

<sup>1</sup> Error limits in pKa estimated from Monte-Carlo analysis are $\pm 0.01$ (see experimental part).
<sup>2</sup> pKa values determined by <sup>31</sup>P-NMR at 298K and by <sup>1</sup>H-<sup>15</sup>N HSQC at 278K

The observation that the apparent acidity of the phosphate groups in the fuzzy complex [23], as compared to those in the isolated disordered domain, increases at high temperature and decreases at 278K is interesting and witness the complexity of the multiple competing interactions present in the fuzzy complex. The importance of electrostatics in fuzzy intermolecular complexes is well recognized [24]

The pH of phosphate buffer decreases by approximately 0.003 pH units per degree [25]. The observed pKa change from 278K to 298K of pT37 and pS43 in pUSH3 (−0.06 units) agrees with this expectation, but the pKa of pS75 moves in the opposite direction. The pKa of phosphate groups in the isolated disordered domain increases with temperature. We can only speculate on the origin of the opposite signs of the pKa temperature coefficients in the fuzzy complex and the isolated disordered region or the opposite trend observed in the pKa changes caused by the presence of the SH3 domain at 278K and 298K. The SH3 domain has a global negative charge, although it contains both positively and negatively charged residues. The fact that the pKa temperature coefficients of p37 and p43 in pUSH3 are similar to those of an isolated phosphate group may suggest that the negative SH3 is competing with the phosphate group for the interaction with the positive charged groups. Also, the decreased apparent acidity observed at the lower temperature, is consistent with the overall expected effect of the phosphate groups being close to the negatively charged SH3 domain.

In the absence of the SH3 domain the interaction between the phosphate group and nearby positively charged groups is unhindered and it may cause the pKa temperature coefficients being opposite in sign to those of an isolated phosphate group. Interestingly the phosphate group of pS75, which shows the strongest basic residue effects on its pKa, is the one showing more extreme temperature coefficients in pUSrc and the only one with a positive coefficient in pUSH3. The opposite effects of the SH3 domain in the pKa of phosphorylated residues at low and high temperature may reflect changes in the compaction of the fuzzy complex or a preferential interaction of the phosphate groups with positively charged residues in the SH3 domain, such as the key arginine residue located in the RT loop of the SH3 domain of Src.

### 2.5. A fast method to identify free or interacting phosphate groups directly from the pH dependency of HSQC spectra

The magnitude of the pH induced chemical shift change provides a direct indication of whether the phosphate group is interacting with distant sites, such as basic residues, or not.

The difference between the high and low pH chemical shifts of the NH of pS75, pS43, and pT37 shows a nearly linear correlation with the pKa shifts experienced by the respective phosphate groups when the interacting basic residue was mutated to alanine (Figure 5).

**Figure 5.**
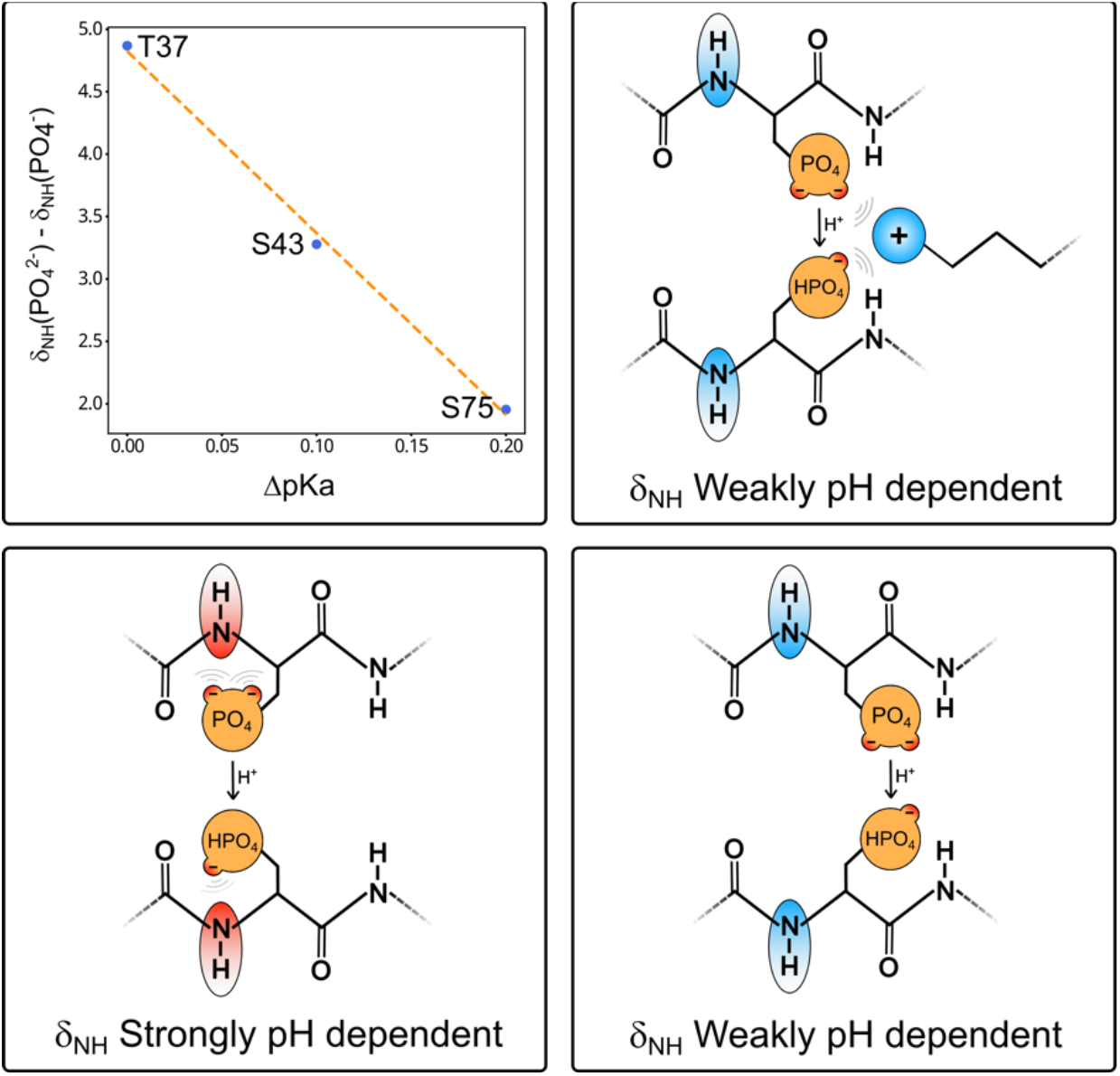
The top left graph shows the linear dependency between the change in pKa induced by the mutation of the nearest basic residue (K40 for K37 and S43; and R78 for S75) and the nitrogen chemical shift difference between the two ionization states of the phosphate group extracted from the fitting of the Henderson-Hasselbalch equation. The cartoons illustrate that the pH dependency of the backbone NH groups of the phosphorylated residue is largest when the phosphate group is directly interacting with the backbone amide of the same residue. The proximity of a basic residue that competes for the interaction with the phosphate group decreases the pH dependency of the NH chemical shifts. Thus, by simply comparing the NH chemical shifts of a phosphorylated residue at pH values well above and below the pKa is enough to determine if the phosphate group is free or is interacting.

The phosphate group of phosphoserine and phosphothreonine can form a hydrogen bond with the NH of the same residue. This interaction is strongly dependent on the ionization state of the phosphate group, so the NH peak experiences large pH-dependent shifts. When the phosphate group participates in a competing external interaction (i.e .with a close basic group), its interaction with the NH group is weakened and the pH-dependent shift is smaller. Thus, recording two HSQC spectra above and below the expected pKa of the phosphate group and measuring the chemical shift change, enables to predict whether the phosphate group is free or is interacting strongly

## 3. Discussion

Phosphorylation is a key post-translational modification through which cell signaling is transmitted. These modifications frequently occur in disordered regions, but their effects are transmitted to the entire protein. The Src N-terminal regulatory element (SNRE) reads the cellular environment through the disordered Unique and SH4 domains to determine the precise outcome of downstream signaling. c-Src is a non-receptor tyrosine protein kinase at the crossroad of many signaling pathways. The Unique domain contains two canonical phosphosites targeted by mitogen activated protein kinases (MAPK). NMR and mass-spectrometry results show that at least two additional sites are phosphorylated in vitro by MAPK1, also known as extracellular regulated kinase (ERK2). All these sites have a basic residue located three-residues upstream or downstream along the sequence.

The pKa of each of the phosphosites has been determined in the isolated disordered region and in the entire SNRE, including the globular SH3 domain, in addition to the disordered domains. Our results clearly show that arginine 78 has a direct interaction with the phosphate group of pS75 that induce a shifts of its pKa of around 0.2 units. Lysine 40 interacts with pS43 resulting in a pKa shift of around 0.1 units. pT37, in spite of having a lysine residue at position i+3 is not affected by the mutation of K40.

Phosphorylation of the Unique domain of c-Src by ERK2 causes smaller but significant pH-dependent chemical shift perturbations in distant residues along the sequence. The pKa of the group causing the observed shifts can be used to identify the specific phosphate group responsible for the perturbation. The effects observed in the guanidinium side chain of R78 and in the intervening residue Q77 provide additional evidence of the direct interaction between pS75 and R78.

In this work we have used point mutations combined with NMR based pKa measurements to identify interactions affecting the phosphorylated residues in the Unique domain. Those are time consuming experiments. However, our data suggests a much simpler approach to identify phosphate groups that have a strong interaction with a basic group: the magnitude of the chemical shift difference of the HN group of the phosphorylated residues correlates inversely with the intensity of the interaction of the phosphate group with neighbor basic residues, as determined by the pKa shift induced when the interacting basic residue is mutated. Thus, by measuring ^1^H-^15^N correlation spectra of a protein with phosphorylated serine or threonine residues at two pH values (*e*.*g*. pH 5.0 and pH 7.5) encompassing the expected pKa of the phosphate group and observing the pH induced shift, one can identify if a particular phosphate group is free or is interacting with a neighbor basic group. ^15^N chemical shifts changes of phosphorylated serine or threonine residues that are not interacting with a basic group have a large pH dependency (of the order of 4-5 ppm) while those having a strongly interacting phosphate show a much smaller dependency (1.5-2 ppm). Thus, a simple preliminary experiment at two pH values provides evidence of the interaction between the phosphate group and basic residues as well as an estimate of the strength of this interaction. Final confirmation will require the complete pKa determination in the wild-type protein as well as basic residue point mutants as described in this paper.

This simple approach has some analogy with the classical identification of intramolecular hydrogen bonds in peptides or proteins through the measurement of amide proton temperature coefficients. In our proposed approach, instead of changing the temperature, we change the pH. Also, similarly to the use of temperature coefficients, the presence of the relevant interactions causes a smaller sensitivity to the external perturbation (low temperature coefficients, small pH dependency of NH chemical shifts). This is due to the competition of the relevant interaction, which has a small dependency on temperature or pH, with other trivial interactions that have a higher sensitivity to the perturbations.

In phosphorylated serine and threonine residues, the phosphate group strongly interacts with the NH group of the phosphorylated residue and the resulting shift in the NH group is strongly pH dependent, as it is directly affected by the ionization state of the phosphate group (Figure 4). When the phosphate group is interacting with a neighbor basic residue, changes in its ionization state have only a weak effect on the NH group of the phosphorylated residue. Thus, the interaction of the phosphate with an external basic group reduces the pH dependency of the NH chemical shifts. It should be clear that this effect applies to phosphorylated serine and threonine residues but not to phosphotyrosine, whose phosphate group cannot interact directly with the backbone NH of the same residue.

The pKa of phosphorylated residues are affected by short- and long-range electrostatic interactions and provide a sensitive probe for the conformational ensemble sampled by disordered domains. pKa measurements at two temperatures, and in the presence or in the absence of the SH3 domain acting as a scaffold of the intramolecular fuzzy complex adopted by the SNRE, reveal a complex scenario. The balance between competing interactions provides a sensitive tests ground to validate calculations of the conformational ensemble adopted by disordered domains [26], especially in the vicinity of SH3 domains [27], although this is beyond the scope of the present article.

## 4. Materials and Methods

### Cloning, prokaryotic gene expression and protein production

Wild type USrc encompassing residues 1-88 of c-Src with a C-terminal streptag and USH3 including residues 1-150 with an N-terminal His_6_-GST fusion had been previously described [23]. The production protocol was modified by using a N-terminal His tagged SUMO fusion protein that enabled affinity purification followed by cleavage with Ubiquitin-like-specific protease 1 (ULP1) that does not leave non-natural residues at the extremes. In the case of the USrc construct, a non-natural tyrosine was introduced at position 86, near the C-terminus to facilitate detection and quantification by UV.

Forward and reverse primers were designed to create point mutants at phosphosites (T37A, S43A, S75A) or at basic residues (K40A, R48A, R78A). QuikChange II XL, PCR site directed mutagenesis kit (Agilent Technologies) was used according to the instruction manual for mutagenesis PCR.

pET28b plasmids encoding His-tagged SUMO at the 5’-end of the corresponding construct were cloned into Omnimax cells and verified by sequencing. The correct plasmids were expressed in *E. coli* BL21 (DE3) pLysS Rosetta cells exploiting kanamycin and chloramphenicol resistencies. Cells were grown in Luria Broth medium for unlabeled samples or minimal M9 medium with ^15^NH_4_Cl or ^15^NH_4_Cl, and [U-^13^C]-glucose for isotopically labelled samples until OD600 reached 0.7. Protein expression was induced by 200 μM isopropyl-β-D-1-thiogalactoside (IPTG) at 25 °C. After induction, cells were centrifuged at 5 000 g for 20 min and the pellet was resuspended in lysis buffer (50 mM Na-Phosphate, 300 mM NaCl, pH 8.0) containing phenyl-methyl-sulfonyl-fluorid (PMSF) and benzamidine protease inhibitors both at 1 mM, and either kept at -80 °C for later use or immediately lysed in the presence of 20 mM imidazole, lysozyme (625 μg/ml) and DNase I (10 μg/ml) by sonication in an ice bath. Cell debris and liquid lysate was separated by 25 minutes at 20 000 rpm. The supernatant was loaded into a NiNTA affinity column (5 ml), washed with lysis buffer, and the desired protein was eluted with 400 mM imidazole. The eluate was dialysed against 50 mM Na-Phosphate, 300 mM NaCl, pH 8.0, 0.5-1.0 mM DTT using 3.5-5 kDa cut-off bag in the presence of ULP1 for at least two hours to remove the excess of imidazole and cleave N-terminal His-tagged SUMO, which was subsequently removed by a second NiNTA column. The flow through was injected in a preparative size exclusion S75 column (approx. 180 ml). In a few instances, anion-exchange chromatography was used before gel filtration. The fractions of gel filtration were checked on 16 % SDS-PAGE and the fractions containing pure protein were concentrated to about 700 μM.

### ERK2 production and phosphorylation reactions

A plasmid encoding a constitutively active, His-tagged form of ERK2 was kindly provided by Prof. Attila Reményi [24] and expressed according to the protocol provided by this group. Briefly, the protein was expressed in *E. coli* BL21 Rosetta (DE3) grown in Terrific Broth medium. Protein induction was carried out with 0.3 mM IPTG when the OD_600_ of the culture reached 0.3-0.4. Following lysis by sonication, the soluble fraction was purified by Nickel affinity using a 5 mL His Trap FF column, eluted with 400 mM imidazole, and further purified by size exclusion using a Sephadex S75 column in 50 mM Tris, 50 mM NaCl, pH 7.4 buffer containing 10% glycerol and 0.01% sodium azide. The purified protein was aliquoted (150 μM, 210 μL), quickly frozen in liquid nitrogen and stored at -80ºC until it was used.

Phosphorylation reactions were carried out at a ERK2 to substrate ratio of 1:50. The reaction conditions were 50 mM phosphate buffer pH 7.4, 5 mM ATP, 10 mM MgCl_2_, 1 mM dithiothreitol (DTT), 3 μM ERK2 and 150 μM substrate. Complete conversion was achieved after three hours at room temperature (approx. 20ºC). USH3 samples were purified by removing ERK2 using a nickel column followed by size exclusion chromatography. The USrc and mutated samples were directly dialyzed against 50 mM citrate buffer (pH 7.2) and concentrated to 70-200 μM.

### Identification of phosphorylation sites by mass-spectrometry

A sample of USH3 phosphorylated by ERK2 gave three distinct bands in a native 8% polyacrylamide gel electrophoresis, corresponding to species with a different number of phosphorylation sites, with no band corresponding to the non-phosphorylated form. The three bands were independently excised from the gel, destained with NH_4_HCO_3_ and acetonitrile, reduced with 10 mM DTT for 45 min at 56ºC and alkylated with 50 mM iodoacetic acid in the dark. Following this treatment, the bands were digested with trypsin (sequencing grade modified trypsin, Proomega Cat#V511) at 37ºC overnight. The digestion was stopped with 5% formic acid and the peptides were eluted with acetonitrile. The sample solution was dried completely in SpeedVac and reconstituted in 20 μL of 3% acetonitrile/1% formic acid aqueous solution for mass spectrometry analysis.

Tryptic digested peptides were trapped with a μ–precolumn 300 μm i.d. x 5 mm Pep-Map100, 5 μm, 100 Å, C18 (Thermo Scientific) separated by a C18 analytical column NanoEase MZ HSS T3 column (75 μm x 250 mm, 1,8 μm, 100 Å) (Waters). The column outlet was directly connected to an Advion TriVersa NanoMate (Advion) fitted on a Orbitrap Fusion Lumos Tribid (Thermo Scientific). Tandem mass-spectrometry (MS/MS) was carried out with a using higher energy collisional dissociation (HCD). We performed a twin database search with Thermo Proteome Discoverer v2.5.0.400 (PD) and MaxQuant v1.6.17.0 (MQ) using the search engines Sequest HT for PD, and Andromeda, for MQ. The databases used were Human and *E. coli* SwissProt release 01/2021 and the common contaminants database as well as a Decoy database to discover the false discovered rate. Search parameters included trypsin enzyme specificity, allowing for two missed cleavage sites, oxidation of methionine, phosphorylation of serine, threonine and tyrosine and acetylation in protein N-terminus as dynamic modifications. The ptmRS node of PD was used to provide a confidence measure for the localization of phosphorylations. Phosphopeptide spectrum matches containing any phosphorylation sites with localization probability < 75% were tagged as ambiguous. For unambiguous sites found with more than 1 peptide spectrum matches (PSM) we computed the phosphorylation ratio (r) for a phosphorylation site from the maximum number of PSM for phosphorylated (N_Phos_) and non-phosphorylated (N_NoPhos_) peptides identified by the Sequest HT and Andromeda nodes. The ratio was computed as r = N_Phos_/(N_Phos_+N_NoPhos_).

### ^31^P NMR measurements and data analysis

NMR measurements were conducted on a Bruker Avance III operating at 9.4 T (400 MHz for ^1^H and 162 MHz for ^31^P) equipped with a sample exchanger. Unless otherwise stated the samples were dissolved in 50 mM citrate buffer and the temperature was 298 K. Triethylphosphate was used as a reference. Spectra were processed and analysed with Topspin 3.4 and MestreLab Research S.L. (analysis)

### pH titration and pK_a_ calculations

For ^31^P-NMR using unlabeled protein, separate samples were adjusted at the desired pH values in the range between 4.8 and 7.2. For isotopically labelled samples, the pH was adjusted between experiments using concentrated HCl or NaOH.

pK_a_ were determined by non-linear fitting to the Henderson-Hasselbalch equation leaving pK_a_ and the high and low pH chemical shifts as adjustable values. The accuracy of the pK_a_ was estimated by Monte Carlo fitting using a collection of 10 artificial data sets generated by adding normal distributed errors to the experimental data points and independently fitting each set to obtain a set of pKa values from which the mean and standard deviations were calculated.

### 2D and 3D NMR experiments

^1^H-^15^N-HSQC experiments were recorded at 278K in a 600 MHz in a Bruker Avance III or a Bruker 800 MHz Avance Neo instruments equipped with TCI cryoprobes. Backbone NH Assignments of double labeled (^13^C-^15^N) USH3 and pUSH3 were confirmed by 3D experiments and transferred to pUSrc. Measuring time was shortened by using best-TROSY versions and a non-uniform sampling strategy (25% NUS) of the following experiments: HNCA, HN(CO)CA, HNCACB, HN(CO)CACB, HNCO, HN(CA)CO and (H)CC(CO)NH-TOCSY at 800 MHz. Cis and trans proline residues were assigned based on the CB and CG chemical shift differences obtained from (H)CC(CO)NH spectra. Side chain assignments of Q77 were based on modified HNCACB experiments. For ε-NH Arg we used the HiSQC approach [28] and connection to the other side chain protons through 2D HCN(CC)H-TOCSY and 3D TOCSY-HSQC ^1^H-^15^N (t_mix_=80 ms) as well as 3D NOESY-HSQC ^1^H-^15^N (t_mix_ = 120 ms). R78 ε-NH showed clear TOCSY transfer along its side chain. R48 gave weaker connections but showed confirming NOEs from HA, HB, HG and HD protons to G49 NH.

## Author Contributions

Conceptualization, M.P.; methodology, M.D-L, M.V., F.C. M.G. ; software, A.F..; investigation, A.L., A.F.; data curation, A.F.; writing, M.P.; supervision, M.P.; project administration, A.F.; funding acquisition, M.P. All authors have read and agreed to the published version of the manuscript.”

## Data Availability Statement

Data are available from the corresponding author.

## Acknowledgments

We thank Attila Reményi for the plasmids to express ERK2, Julie Forman-Kay and Rhea Hudson, for pointing out the use of ERK2 to phosphorylate c-Src and for sharing plasmids and protocols, Dr. Christopher D. Lima and Cornell University for the license to use cleavable SUMO fusion proteins, Gianluca Arauz-Garofalo for bioinformatic support in mass-spectrometry. Mass spectrometry/Proteomics was a member of Proteored, PRB3-ISCIII and is granted to set up and develop a Proteomics Platform for Biomedical and Translational Research in the framework of the 2014-2020 ERDF Operational Programme in Catalonia. the authors thankfully acknowledge the NMR resources and the technical support provided by the LRB (Laboratorio de RMN de Barcelona) of the Spanish ICTS Red de Laboratorios de RMN de Biomoléculas (R-LRB.

